# Sex differences and similarities in the neuroimmune response to central administration of poly I:C

**DOI:** 10.1101/2020.11.23.394742

**Authors:** Caitlin K. Posillico, Rosa E. Garcia-Hernandez, Natalie C. Tronson

**Affiliations:** Psychology Department, Ann Arbor 48109; University of Michigan, Ann Arbor 48109

**Keywords:** poly I:C, sex differences, neuroimmune, cytokines, hippocampus, chemokines, male, female, interferon, mouse

## Abstract

**Background:** The neuroimmune system is required for normal neural processes, including modulation of cognition, emotion, and adaptive behaviors. Aberrant neuroimmune activation is associated with dysregulation of memory and emotion, though the precise mechanisms at play are complex and highly context dependent. Sex differences neuroimmune activation and function further complicate our understanding of its roles in cognitive and affective regulation.

**Methods:** Here, we characterized the physiological sickness and inflammatory response of the hippocampus following intracerebroventricular (ICV) administration of a synthetic viral mimic, polyinosinic:polycytidylic acid (poly I:C), in both male and female C57Bl/6 mice.

**Results:** We observed that poly I:C induced weight loss, fever, and elevations of cytokine and chemokines in the hippocampus of both sexes. Specifically, we found transient increases in gene expression and protein levels of IL-1*α*, IL-1*β*, IL-4, IL-6, TNF*α*, CCL2, and CXCL10, where males showed a greater magnitude of response compared with females. Only males showed increased IFN*α* and IFN*γ* in response to poly I:C, whereas both males and females exhibited elevations of IFN*β*, demonstrating a specific sex difference in the anti-viral response in the hippocampus.

**Conclusion:** Our data suggest that type I interferons are one potential node mediating sex-specific cytokine responses and neuroimmune effects on cognition. Together, these findings highlight the importance of using both males and females and analyzing a broad set of inflammatory markers in order to identify the precise, sex-specific roles for neuroimmune dysregulation in neurological diseases and disorders.

## 1. BACKGROUND

The neuroimmune system is responsible for surveying the microenvironment and responding to illness, injury, and infection. Importantly, it is also required to carry out normal neural processes (1–3), thus making it well-placed to modify or disrupt learning and memory. In fact, dysregulated immune signaling has been implicated in disorders of affect and cognition, many of which show sex-biases in prevalence and outcomes (4–6). There is significant evidence for sex differences in immune responses in the periphery (7, 8), but there is limited literature on similar findings in adult brains. Knowing whether such sex differences occur in similar magnitudes and directions in the neuroimmune system is important for understanding exactly how neuroimmune dysregulation impacts cognition, and contributes to psychiatric and neurological disorders, in both sexes.

Illness, injury, or aseptic triggers of the innate immune system – either bacterial endotoxins (e.g., lipopolysaccharide, LPS) or viral mimics (e.g., polyinosinic:polycytidylic acid, poly I:C) – cause activation of neuroimmune cells, including microglia and astrocytes, and rapid production of cytokines in the brain (9, 10). Due to key roles in peripheral inflammation, the cytokines IL-1β (11–13), IL-6 (14–16), and TNFα (17–19) have been the focus of much of the research of neuroimmune function (20). More recently, other cytokines, including interferons (4, 21), CCL2 (22, 23), and CXCL10 (21,24,25) also play critical roles in modulation of behavior, cognition, and affective states, suggesting that many cytokines play important roles in these processes.

Sex differences in immune and neuroimmune activity have also been reported in various contexts. Females have a greater peripheral immune response compared with males (7). In contrast, neuroimmune cells *in vitro*, including astrocytes derived from male cortical tissue, have a significantly greater reaction to inflammatory insults compared with female-derived cells (26, 27). We have identified sex differences in the magnitude, time course, and pattern of cytokines activated in the hippocampus following peripheral LPS (28), and in the long-lasting impact of LPS on hippocampal function (29). Thus, sex differences in neuroimmune responses specifically may be a contributing factor to sex differences in neural and cognitive processes and disorders.

Despite incredible advances in psychoneuroimmunology over the past decade, there are critical gaps in our knowledge that preclude a holistic understanding of neuroimmune function and its impacts on cognition and disease. First, with some notable exceptions (22,30,31), studies have typically focused on a few inflammatory cytokines (e.g., IL-1*β*, IL-6, and TNF*α*) critical for neuroimmune activation and its effects on cognition. Yet more recently, it has become clear that the massive, coordinated cytokine response observed in the periphery also occurs in the CNS (28, 32). The roles played by other cytokines, and the similarities and differences of this response in males versus females are yet to be defined (20). Second, the bulk of studies aimed at understanding neuroimmune activation and its behavioral sequelae have used the gram-negative bacterial shell and toll-like receptor 4 (TLR4) agonist LPS. Nevertheless, viral illnesses also trigger changes in behavior, cognition, and emotional states, and significant sex differences have been observed in the context of viral infections (7,8,33) – an issue that has been propelled to the forefront during the current COVID-19 pandemic (34, 35). Given that viruses act through distinct toll-like receptors, their impact is likely mediated by a different, albeit overlapping, pattern of cytokine activation compared with LPS or bacterial triggers. Third, due to its relevance for disease states, many *in vivo* studies of neuroimmune function use a peripheral immune challenge. Here, neuroimmune activation is primarily driven by peripheral immune signals that infiltrate the brain (36). This complicates the interpretation of whether sex differences in cytokine levels observed in the brain are due to indirect effects based on sex differences in peripheral immune response, or to direct effect of sex differences in neuroimmune function.

In this study, we aimed to identify a broad set of inflammatory cytokines induced in the hippocampus by direct neuroimmune stimulation *via* central administration of poly I:C in both males and females. Elevation of hippocampal cytokines, in particular, is associated with both disruption of memory processes (1,11,13,37–40) and increased depression-like behaviors (41, 42). Here, we demonstrate that poly I:C induces fever, weight loss, and changes in mRNA expression and protein levels of cytokines, chemokines, and markers of glial activation across a 24-hour period in both sexes. Notably, only IFNα and IFNɣ showed male-specific patterns of activation after central poly I:C administration, and many cytokines and chemokines showed a greater magnitude increase in males compared with females. Whether these sex differences in neuroimmune activation contribute to sex differences in modulation of cognition and affect and subsequent prevalence of memory- and mood-related diseases and disorders is an important area of research for our ongoing studies.

## 2. METHODS

### 2.1. Animals

A total of 99 male and female 8-9 week-old C57BL/6N mice from Envigo (Indianapolis, IN) were used in these experiments. For all experiments, mice were individually housed in standard mouse cages with *ad libitum* access to food and water in a room with maintained temperature and pressure under a 12:12-hour light:dark cycle. All mice had at least one week of acclimation to the colony room prior to any manipulations. All protocols were approved by the University of Michigan Institutional Animal Care and Use Committee (IACUC).

### 2.2. Stereotaxic Surgeries

Bilateral guide cannulae (PlasticsOne, Roanoke, VA) targeting the lateral ventricles were implanted using standard stereotaxic methods (KOPF, Tujunga, CA) at the following coordinates relative to Bregma: medial-lateral: +/- 1.00mm, anterior-posterior: 0.30mm, dorsal-ventral: -2.50mm. Animals were administered a pre-surgical analgesic (5mg/kg Carprofen, subcutaneous) and anesthetized for surgery using an intraperitoneal injection of 250mg/kg of Avertin (2,2,2-tribromoethanol) which maintained a surgical plane of anesthesia for the duration of the craniotomy. Bilateral holes were drilled into the skull at the above coordinates, and guide cannulae were implanted using dental cement. Animals were given a second dose of Carprofen (5mg/kg, subcutaneous) 24 hours after surgery to maintain a total of 48 hours of analgesia. Mice were monitored daily for 10 days post-operative and were given at least 2 weeks to recover from surgery prior to use in experiments.

### 2.3. Poly I:C Administration

Polyinosinic:polycytidylic acid (poly I:C; Cat. No. P1530; Sigma-Aldrich, St. Louis, MO) was prepared according to the manufacturer’s instructions and sterile-filtered using a 0.22μm filter prior to administration. For intracerebroventricular (ICV) administration, we infused 20μg of poly I:C (2μL of 10μg/μL poly I:C) (43) or an equal volume of 0.9% sterile saline *via* the implanted guide cannula under brief isoflurane anesthesia.

### 2.4. Sickness Behavior Assessment

To determine whether poly I:C increased physiological measures of sickness in males and females, body weights and rectal temperatures (RET-3; Physitemp, Clifton, NJ) were assessed at 2, 4, 6, 12, 24, and 48 hours following ICV administration of poly I:C (*n* = 10 male; *n* = 9 female) or sterile saline (*n* = 10 male; *n* = 8 female; Figure 2A). Visual and behavioral measures of sickness (piloerections, squinted eyes, hunched posture, and low responsivity) were assessed throughout (44). No changes in overt sickness behaviors were observed for any experiment (data not shown).

#### 2.4.1. Statistical Analysis of Sickness Behaviors

Analysis of body weight and temperature changes in response to poly I:C was completed using a mixed repeated-measures ANOVA, using time post-infusion as the within- subjects factor and treatment and sex as the between-subjects factors with Greenhouse-Geisser corrections for sphericity. Significant main effects and interactions (p < 0.05) were followed up using post-hoc tests with Bonferroni corrections for multiple comparisons, and effect sizes were calculated using the partial eta squared method. Any outliers were identified as samples outside the range of 2 standard deviations from the group mean.

### 2.5. Characterization of the Acute Neuroimmune Response

We used RNA and protein endpoints to examine induction of cytokines and glial activation markers in the hippocampus. Males and females were treated with either poly I:C (*n* = 22 male; *n* = 24 female) or sterile saline (*n* = 8/sex) and brains were collected 0.5 hours (*n* = 5 male; *n* = 6 female), 2 hours (*n* = 6/sex), 4 hours (*n* = 5 male; *n* = 6 female), and 24 (*n* = 6/sex) hours later. All animals were transcardially perfused with 0.1M phosphate buffer to remove circulating blood from the brain. Both hemispheres of dorsal hippocampus tissue were collected in separate RNase-/DNase-free, sterile microcentrifuge tubes and immediately flash frozen. All samples were stored at -80°C before tissue processing.

#### 2.5.1. Quantitative Real-Time PCR

One hemisphere of dorsal hippocampal tissue per mouse was processed for gene expression analysis using quantitative real-time PCR (qPCR). Frozen samples were homogenized, and messenger RNA (mRNA) was extracted (PureLink RNA Mini Kit; Cat. No. 12183020; Invitrogen, Carlsbad, CA) under sterile, RNase-free conditions. RNA quality was assessed using gel electrophoresis, and UV spectroscopy was used to assess RNA purity (A260/280 > 1.80) and quantity (BioSpectrometer Basic; Eppendorf, Hamburg, Germany). Any genomic DNA in the sample was removed using DNase treatment, and 800ng of cDNA was synthesized from each mRNA sample (QuantiTect Reverse Transcriptase Kit; Cat. No. 205314; Qiagen, Hilden, Germany). Any samples that did not have a high enough concentration of RNA to make 800ng of cDNA were removed from further analyses (*n* = 3 male; *n* = 5 female). Relative gene expression was measured using Power SYBR Green PCR Master Mix (Cat. No. 4368702; Applied Biosystems, Foster City, CA) in 10μL reactions (ABI 7500 real-time PCR system; Cat. No. 4351105; Applied Biosystems).

We measured expression of four commonly-used housekeeping genes: *18s*, *gapdh*, *hprt1*, and *rplp0* (all QuantiTect Primer Assays: *18s* Cat. No. QT02448082, *gapdh* Cat. No. QT01658692, *hprt1* Cat. No. QT00166768, *rplp0* Cat. No. QT00249375; Qiagen). We analyzed the relative expression of the following genes of interest: *ccl2*, *cd11b*, *cxcl10*, *gfap*, *ifnα*, *ifnβ*, *ifnγ*, *il-1α*, *il-1β*, *il-6*, *il-10*, and *tnfα*. The gene primer for *il-1α* was a QuantiTect Primer Assay (Cat. No. QT00113505; Qiagen). The sequences for the remaining gene primers can be found in Table 1 and were ordered through Integrated DNA Technologies and diluted to 0.13 μM to be used for PCR. All Qiagen primers were diluted as per the manufacturer’s instruction.

**Table 1.**
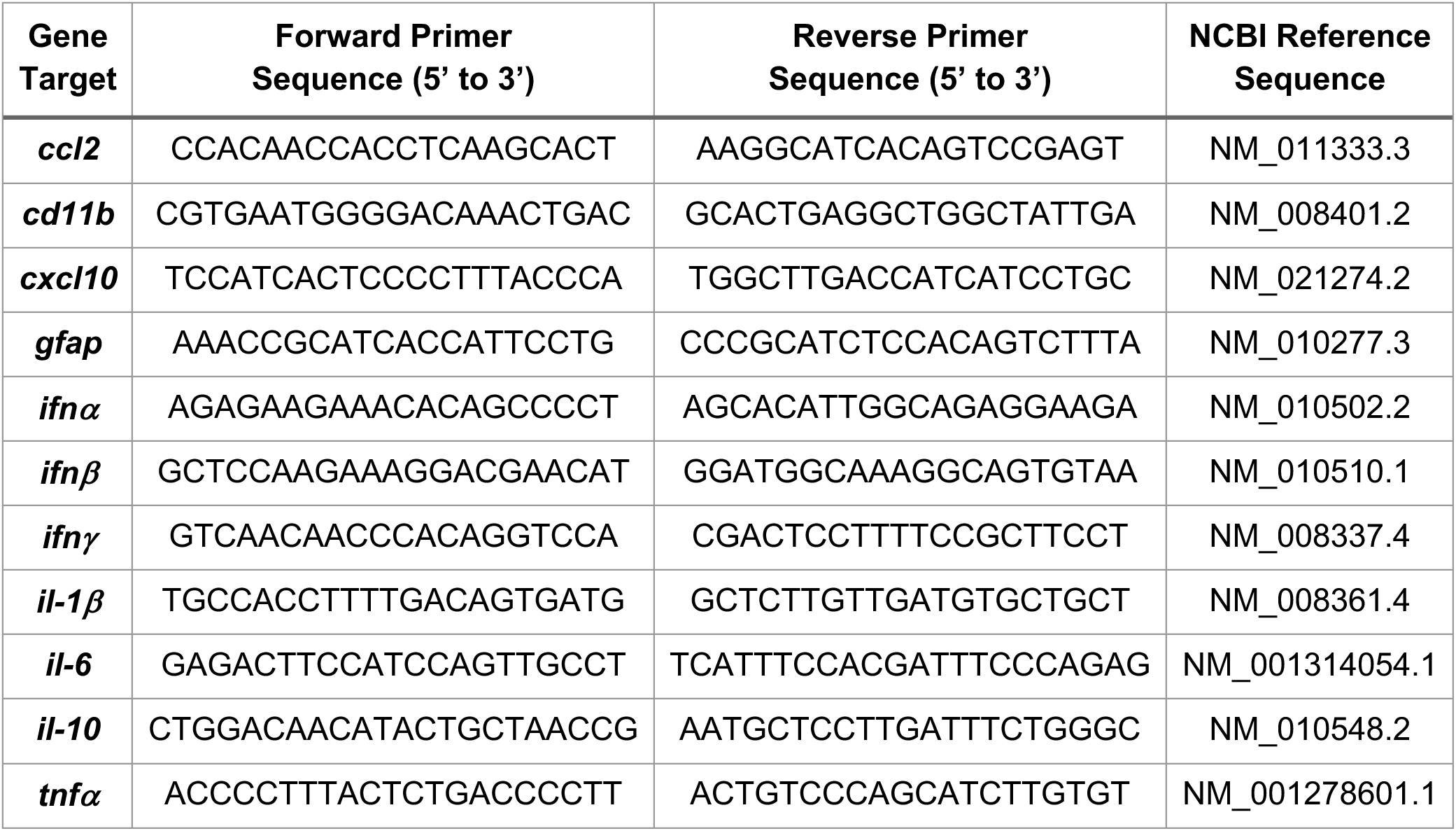
Primer sequences used for real-time PCR.

##### 2.5.1.1. Housekeeping Gene Stability Analysis

To control for the transcriptional activity of the samples being analyzed, we confirmed the stability of four housekeeping genes (*18s*, *gapdh*, *hprt1*, and *rplp0*). While many studies use common housekeeping genes such as GAPDH or HPRT1, it is less common for authors to report that their chosen housekeeping gene is indeed stable across experimental groups or tissues prior to use in analyses. Thus, we confirmed the stability of our housekeeping genes using a combination of four techniques to ensure the most reliable quantification of gene expression in our studies. First, we assessed the variability of the candidate genes by measuring the standard deviation of the raw quantification cycle (Cq) values from all samples (Figure 1A). We found that *18s* had the largest standard deviation of Cq values (1.540), followed by *gapdh* (0.527), *rplp0* (0.225), and *hprt1* (0.151; Figure 1B). By this approach, *rplp0* and *hprt1* showed the greatest stability compared to *18s* and *gapdh*, with *hprt1* exhibiting the lowest variability.

**Figure 1.**
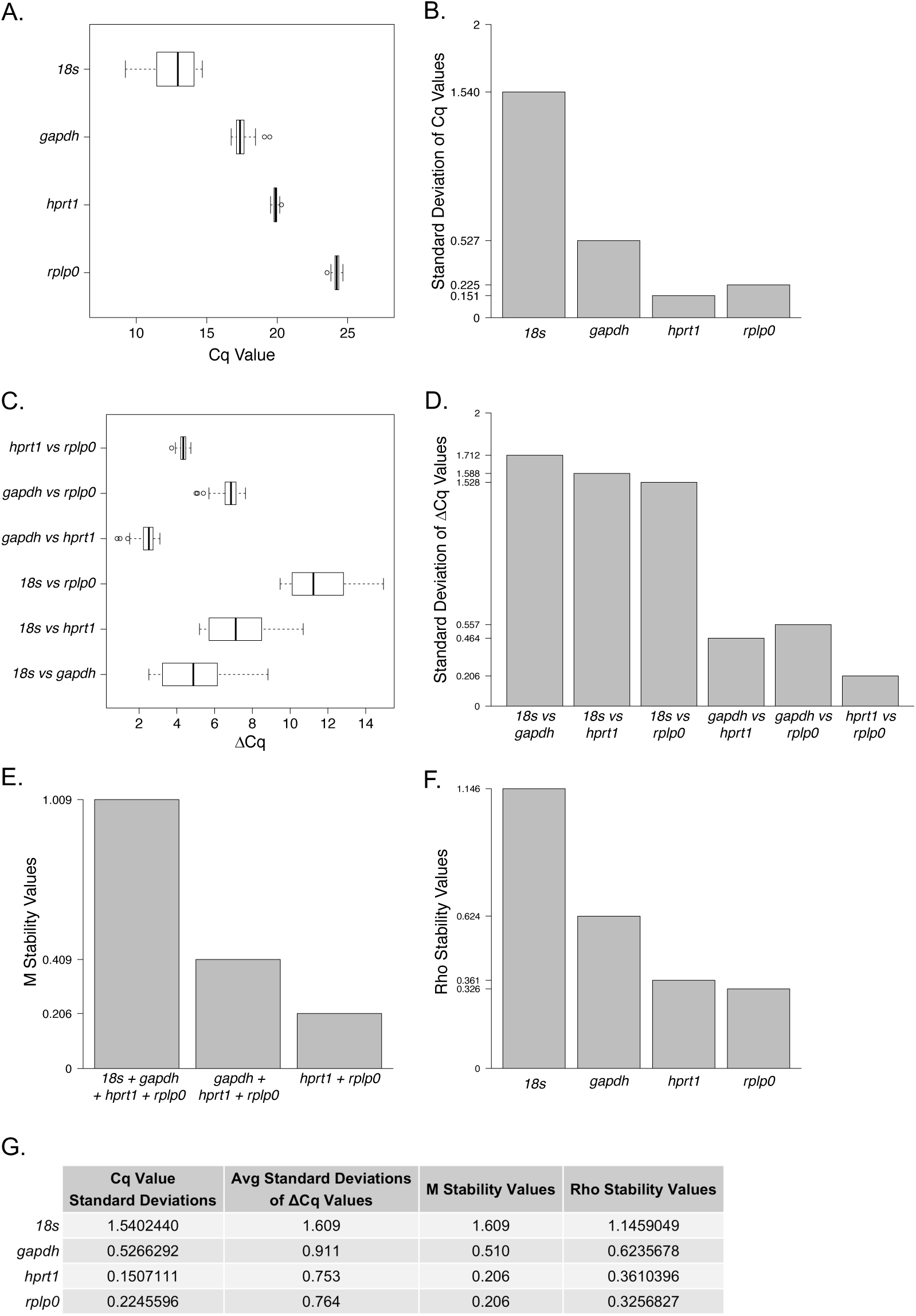
Housekeeping gene stability analysis. **(A)** Distribution of the quantification cycles (Cq) for housekeeping genes *18s*, *gapdh*, *hprt1*, and *rplp0*, with **(B)** associated standard deviations. **(C)** Distribution of the difference of Cq values (*Δ*Cq) between pairs of housekeeping genes, and **(D)** the associated standard deviations. **(E)** Stability values calculated using gene ratio method by Vandesompele et al., 2002, which uses stepwise elimination of lowest stability (highest *M* value) to rank gene stability. **(F)** Stability values calculated using a model-based approach by Andersen et al., 2004 which measures expression variation such that highest stability results in the lowest Rho value. **(G)** Summary of results from each of the four methods of housekeeping gene stability are shown.

Second, we employed a comparative *Δ*Cq approach in which the standard deviations of the differences in Cq values (*Δ*Cqs) between all possible pairs of candidate genes were compared (45) (Figure 1C). From highest to lowest variability, the genes ranked as follows: *18s* (1.609 average standard deviation), *gapdh* (0.911), *rplp0* (0.764), and *hprt1* (0.753). Again, this method indicated that the most variable genes were *18s* and *gapdh* while the most stable genes were *rplp0* and *hprt1*, and this is most apparent when considering the lowest *Δ*Cq standard deviation from this method was from the *rplp0* and *hprt1* comparison at 0.206 (Figure 1D).

The third method we employed was that developed by Vandesompele and colleagues, which calculated the average pairwise variation of one candidate gene with all other candidate genes (46). We used R packages ReadqPCR and NormqPCR (47) to calculate *M* stability values, as depicted in Figure 1E. Consistent with the previous methods, *hprt1* and *rplp0* were the most stable of the candidate genes, with the lowest pairwise variability, *M* value, of 0.206.

Fourth, and last, we used a model-based stability analysis approach developed by Andersen et al., an algorithm called NormFinder (v5) (48). This method protects against identifying two genes via the pairwise approach that might be misinterpreted as being the most stable if they are coregulated. Using this method, again, *hprt1* and *rplp0* were found to be the most stable genes with the lowest expression stability values (Figure 1F). However, NormFinder resulted in *rplp0* having the lowest stability value of 0.326, indicating that the model-based approach identified *rplp0* as the most stable gene.

Together, these methods identified the two most stable candidate housekeeping genes as *hprt1* and *rplp0*. Vandesompele et al. (46) posits that using the geometric mean of multiple housekeeping genes results in more accurate expression levels of genes of interest. We calculated the geometric mean of the Cq values from *hprt1* and *rplp0* to be used in the 2^-ΔΔCq^ method for calculations of relative expression for our target genes.

##### 2.5.1.2. Statistical Analysis of mRNA Gene Expression

For each PCR reaction, the quantification cycle (Cq) was determined, and the 2^−ΔΔCq^ method was used to calculate the relative gene expression of each gene. Any samples with abnormal amplification curves, melt curves, and/or melt peaks across replicates were removed from analyses (*n* = 1/sex). Any outliers were identified as samples outside the range of 2 standard deviations from the group mean and excluded from analyses.

Baseline sex differences in relative gene expression (qPCR) were assessed by evaluating the male and female saline-treated groups. To directly and meaningfully compare these two groups in the PCR analysis, the male saline-treated group was normalized to the female saline-treated group and analyzed using independent, two-sample t-tests.

To appropriately analyze sex differences in relative gene expression (qPCR) across the 24-hour time course, we normalized each group to its respective same-sex saline-treated group to control for any sex differences in gene expression at baseline and used two-way ANOVA tests using treatment and sex as factors. Significant main effects and interactions (*p* < 0.05) were followed up using post-hoc tests with Bonferroni corrections for multiple comparisons, and effect sizes were calculated using the partial eta squared method.

#### 2.5.2. Multiplex Assays

The second hemispheres of dorsal hippocampal tissue were processed as previously described using low-detergent RIPA buffer sonication (28). Milliplex magnetic bead panel assays (CCL2, CXCL10, IFN*γ*, IL-1*α*, IL-1*β*, IL-2, IL-4, IL-6, and IL-10; Millipore Sigma, Burlington, MA) were used as per manufacturer’s instructions. Cytokine concentrations were calculated as pg/mg of hippocampal tissue via Luminex software. Only samples that showed readable bead counts according to the Luminex software were included in the analyses.

##### 2.5.2.1. Statistical Analysis of Protein Levels

Baseline sex differences in protein levels from multiplex assays were analyzed with independent, two-sample t-tests comparing the saline-treated groups. To analyze changes in protein levels from poly I:C across the 24-hour time frame, we used two-way ANOVA tests using treatment and sex as factors. Significant main effects and interactions (*p* < 0.05) were followed up using post-hoc tests with Bonferroni corrections for multiple comparisons, and effect sizes were calculated using the partial eta squared method. Any outliers were identified as samples outside the range of 2 standard deviations from the group mean and excluded from analyses.

### 2.6. Data Visualization and Statistical Software

Data visualization and statistical analyses were completed using R 3.6.2 (R Core Team, 2019) with the following packages: dplyr (v0.8.5; (49)), tidyr (v1.0.2; (50)), rstatix (v0.5.0; (51)), DescTools (v0.99.34; (52)), sjstats (v0.17.9; (53)), ReadqPCR and NormqPCR (47), ggplot2 (54), gridExtra (v2.3; (55)), pheatmap (v1.0.12; (56)), and viridis (v0.5.1; (57)).

## 3. RESULTS

### 3.1. Central administration of poly I:C induces physiological sickness responses

Both females and males showed physiological responses to poly I:C. Whereas both saline- and poly I:C-treated animals showed changes in weight across the 48-hour period (Figure 2B, main effect of Time: *F*(3.13, 96.92) = 28.899, *p* < 0.001, η^2^*_p_* = 0.482), poly I:C caused weight loss in both sexes (main effect of Treatment: *F*(1, 31) = 8.781, *p* = 0.006, η^2^*_p_* = 0.221; trend towards a Time x Treatment interaction: *F*(3.13, 96.92) = 2.476, *p* = 0.064, η^2^*_p_* = 0.074). Specifically, males and females treated with poly I:C lost significantly more weight than the saline-treated animals at the 12- (*p* = 0.004) and 24-hour (*p* = 0.022) time points. By 48 hours post-treatment, the weights of poly I:C-treated animals had recovered and were no longer different from those of saline-treated animals (*p* = 1.00).

**Figure 2.**
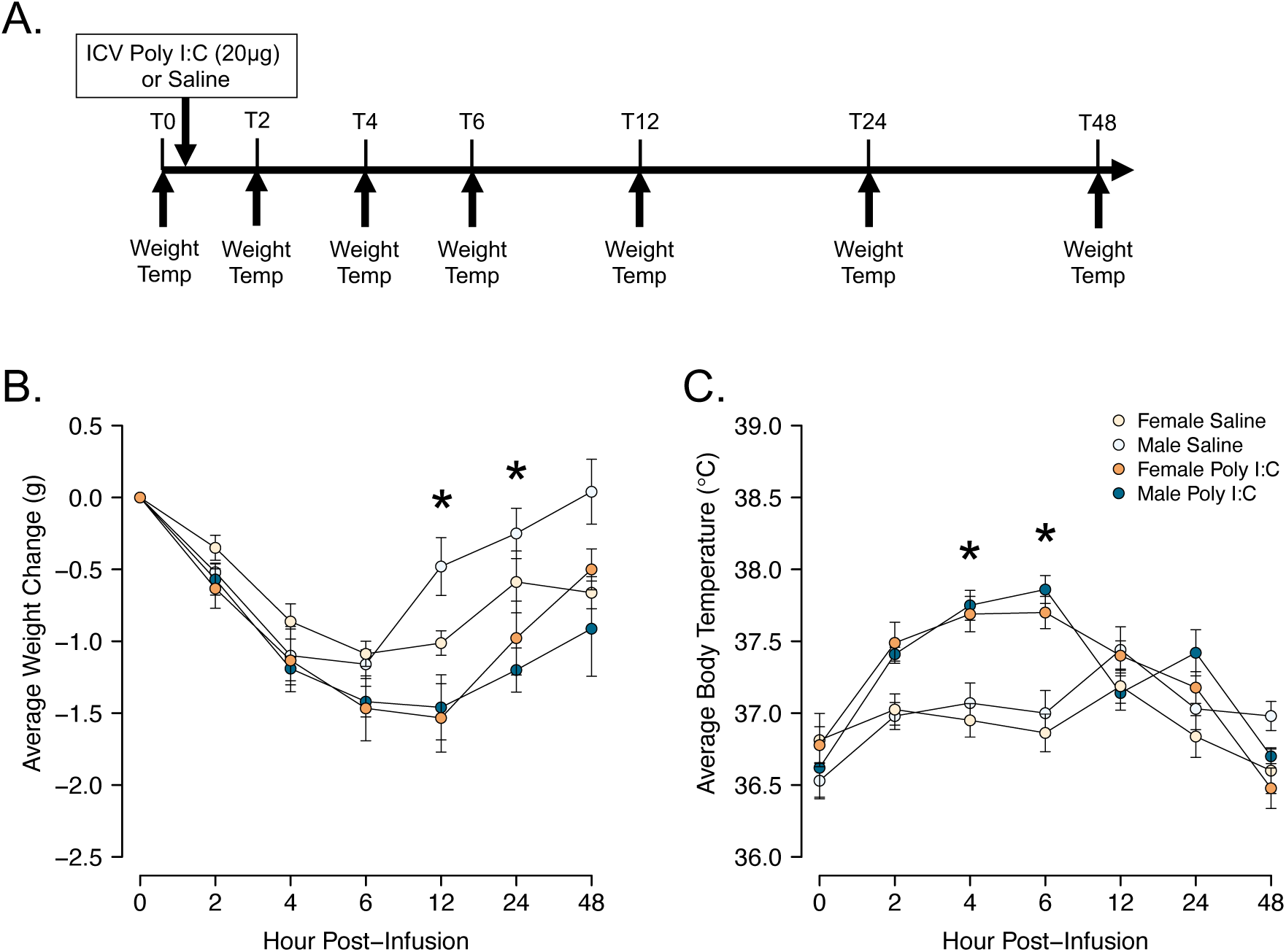
Analysis of sickness behaviors following poly I:C administration. **(A)** Timeline of body weight and temperature measurements following poly I:C or sterile saline administration. **(B)** Average weight change from baseline (Time = 0 hour) prior to treatment. **(C)** Average body temperature as measured via rectal thermometer. Analyzed using mixed repeated-measures ANOVA. **\***: *p* < 0.05 poly I:C- vs saline-treated groups.

In both males and females, poly I:C caused significant increases in body temperature relative to the saline-treated group (Figure 2C; main effect of Treatment: *F*(1, 31) = 23.759, *p* < 0.001, η^2^*_p_* = 0.434; Time x Treatment interaction: *F*(4.6, 142.62) = 11.635, *p* < 0.001, η^2^*_p_* = 0.273). Post-hoc tests revealed that body temperature began to increase 2 hours following poly I:C (*p* = 0.068), remained elevated at the 4- (*p* < 0.001) and 6-hour (*p* < 0.001) time points, and recovered to saline-treated body temperatures by 12 hours post treatment (all *p* = 1.00).

### 3.2. Baseline sex differences in mRNA expression and protein levels of select hippocampal immune molecules

#### 3.2.1. Interleukins

Select immune signaling markers showed baseline sex differences. mRNA expression of interleukin *(il)-1α* exhibited a trend towards a sex difference in baseline expression (Figure 3C1; *t*(12) = 2.006, *p* = 0.068), and *il-6* showed a significant sex difference (Figure 3E1; *t*(11) = 3.079, *p* = 0.01, 95% CI [0.182, 1.062]) where females showed greater mRNA expression than males. These results did not hold true for protein levels of IL-1*α* or IL-6. Rather, multiplex data showed that protein levels of IL-1*β* and IL-10 were significantly different between saline-treated males and females (Figures 4D and 4F, respectively; IL-1*β*: *t*(13) = 4.275, *p* = 0.001, 95% CI [5.682, 17.291]; IL-10: *t*(13) = 2.236, *p* = 0.044, 95% CI [0.672, 39.314]). Again, for these cytokines, females showed greater protein levels at baseline compared with males.

**Figure 3.**
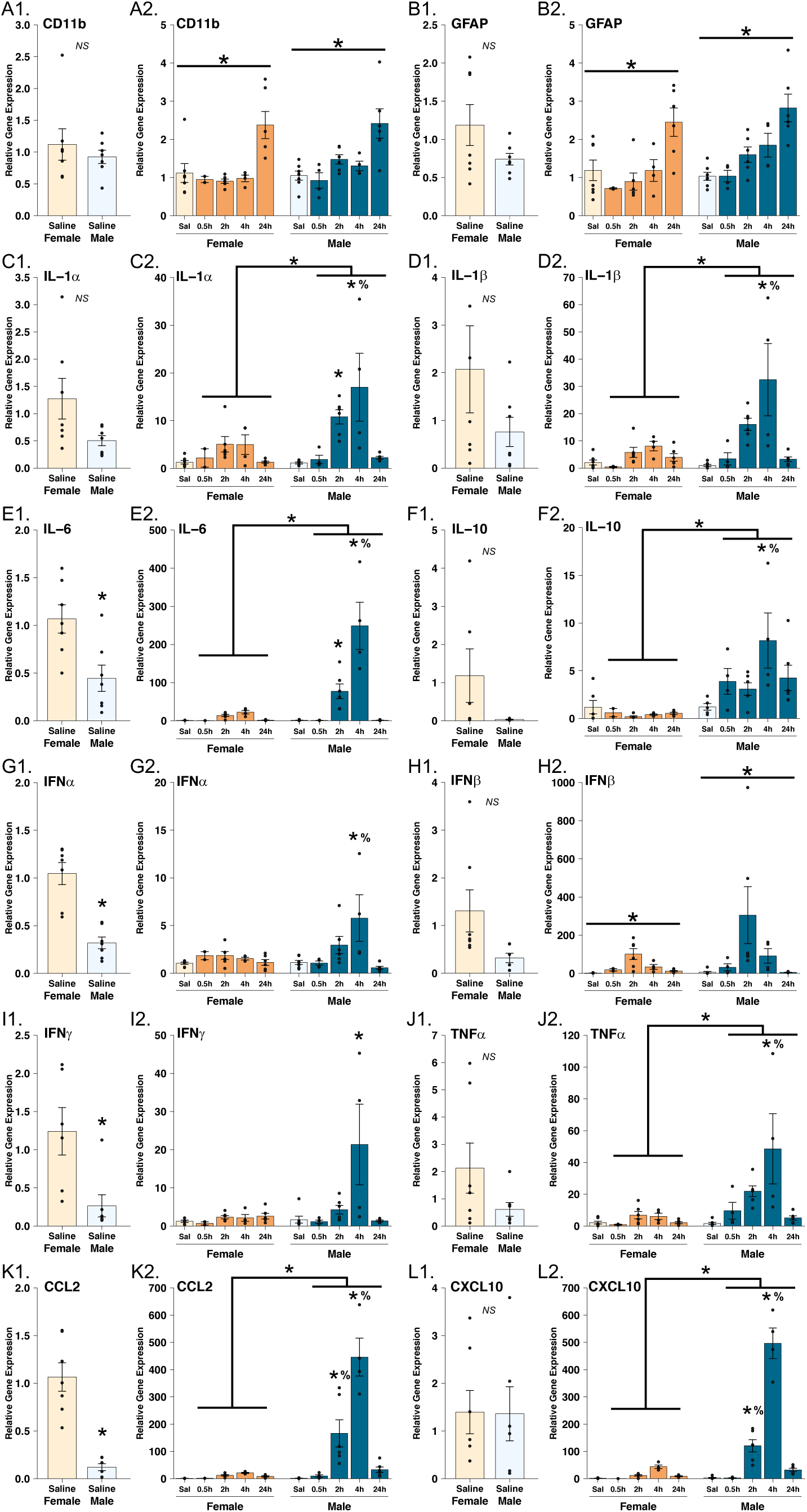
mRNA gene expression of cytokines, chemokines, and markers of glial activation in the hippocampus. Baseline gene expression was measured by normalizing the male saline-treated group to the female saline-treated group and analyzed using independent, two-sample t-tests. Baseline expression of **(A1)** CD11b, **(B1)** GFAP, **(C1)** IL-1*α*, **(D1)** IL-1*β*, **(E1)** IL-6, **(F1)** IL-10, **(G1)** IFN*α*, **(H1)** IFN*β*, **(I1)** IFN*γ*, **(J1)** TNF*α*, **(K1)** CCL2, and **(L1)** CXCL10 are shown. Gene expression changes following poly I:C treatment were calculated by normalizing time points after poly I:C treatment to the saline-treated groups within sex to eliminate confounding variables of baseline sex differences. Gene expression was analyzed using two-way ANOVA tests for **(A2)** CD11b, **(B2)** GFAP, **(C2)** IL-1*α*, **(D2)** IL-1*β*, **(E2)** IL-6, **(F2)** IL-10, **(G2)** IFN*α*, **(H2)** IFN*β*, **(I2)** IFN*γ*, **(J2)** TNF*α*, **(K2)** CCL2, and **(L2)** CXCL10. **\*** above a bracket covering both sexes indicates a significant main effect of sex (*p* < 0.05); * above a horizontal line covering just one sex indicates a significant main effect of treatment (*p* < 0.0.5); **\*** above a single bar indicates a significant post-hoc test (*p* < 0.05) vs the saline-treated group within sex; **%** above a single bar indicates a significant post-hoc test (*p* < 0.05) vs females at the same time point.

**Figure 4.**
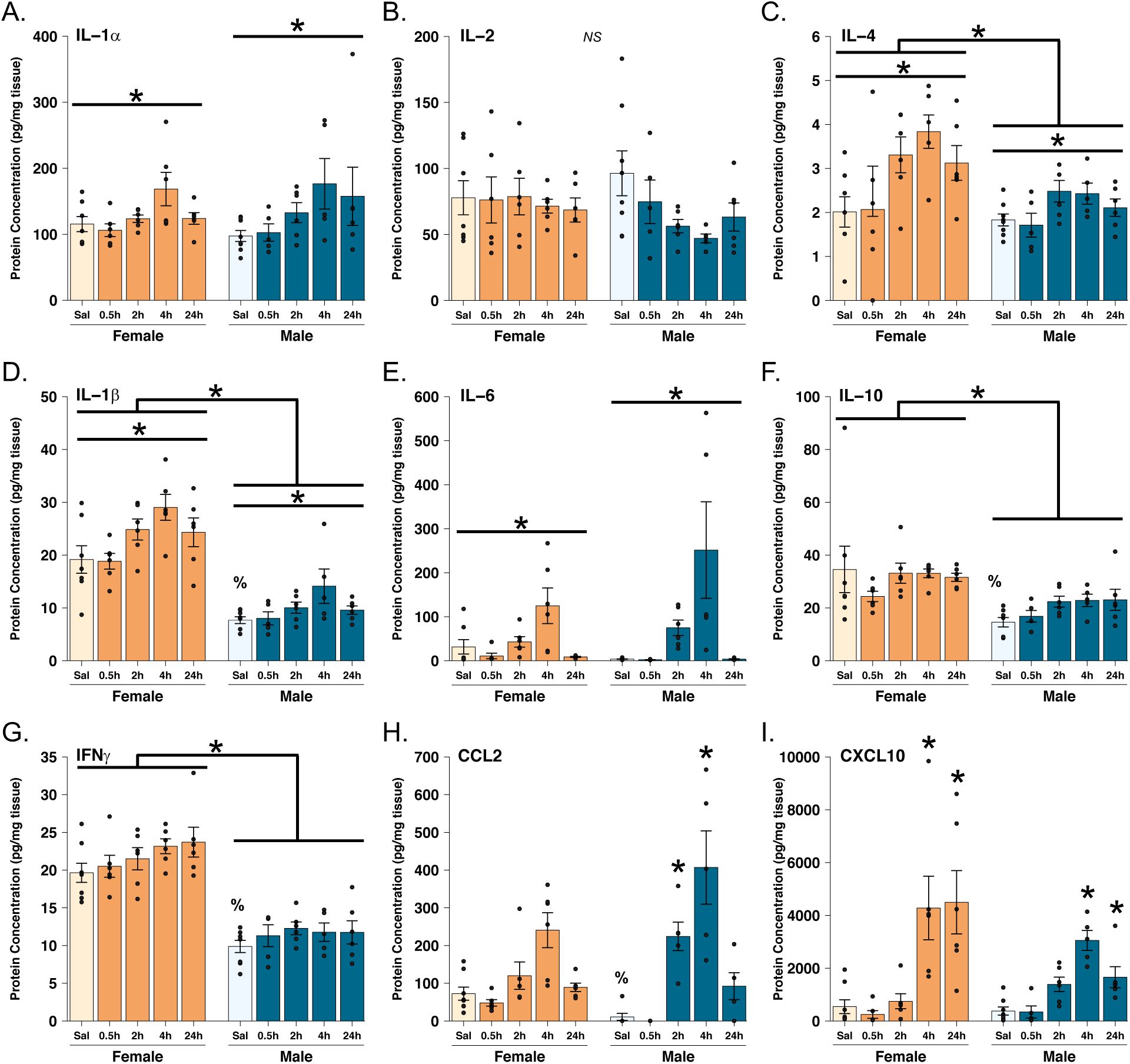
Protein levels of cytokines and chemokines in the hippocampus. Protein concentration (pg/mg) of **(A)** IL-1*α*, **(B)** IL-2, **(C)** IL-4, **(D)** IL-1*β*, **(E)** IL-6, **(F)** IL-10, **(G)** IFN*γ*, **(H)** CCL2, and **(I)** CXCL10 are shown. Two-way ANOVA tests were used to analyze these data. **\*** above a bracket covering both sexes indicates a significant main effect of sex (*p* < 0.05); * above a horizontal line covering just one sex indicates a significant main effect of treatment (*p* < 0.0.5); **\*** above a single bar indicates a significant post-hoc test (*p* < 0.05) vs the saline-treated group within sex; **%** above a single bar indicates a significant post-hoc test (*p* < 0.05) vs females at the same time point.

#### 3.2.2. Interferons

Two interferons (IFN), *ifnα* and *ifnγ*, also showed baseline sex differences in relative mRNA expression (Figures 3G1 and 3I1: IFN*α*: *t*(12) = 5.546, *p* = 0.0001, 95% CI [0.441, 1.01]; IFN*γ*: *t*(11) = 2.995, *p* = 0.012, 95% CI [0.259, 1.694]). Likewise, protein levels of IFN*γ* exhibited a sex difference in saline-treated groups (Figure 4G; *t*(14) = 6.475, *p* < 0.001, 95% CI [6.534, 13.006]). For both mRNA expression and protein levels, female levels were greater than males.

#### 3.2.3. Chemokines

Expression of CC chemokine *ccl2* similarly followed this pattern at baseline (Figure 3K1; *t*(12) = 3.287, *p* = 0.006, 95% CI [0.259, 1.279]), and this held true for protein levels of CCL2 as well (Figure 4H; *t*(12) = 2.751, *p* = 0.018, 95% CI [12.798, 110.318]). Interestingly, all genes with sex differences in saline-treated animals showed that both female baseline mRNA expression and protein levels were at least twice as high as that of males.

#### 3.2.4. Glial activation markers and other cytokines did not show baseline sex differences

It is equally important to note the markers that did not show sex differences at baseline. Both the microglial activation marker *cd11b* and the astrocyte activation marker *gfap* showed equivalent levels of gene expression in the saline-treated groups in both sexes (Figures 3A1 and 3B1, respectively; *cd11b*: *t*(12) = 0.723, *p* = 0.483; *gfap*: *t*(12) = 1.603, *p* = 0.135). Other commonly examined inflammatory cytokines, including IL-2, IL-4, and tumor necrosis factor (TNF)*α* similarly showed no sex difference at baseline (IL-2 p = 0.420; IL-4 p = 0.633; TNF*α* = 0.139). No other sex differences in mRNA gene expression or protein levels were observed under baseline conditions.

### 3.3. Central poly I:C causes greater increases in male mRNA expression of hippocampal cytokines compared with females

#### 3.3.1. Glial Activation Markers

Poly I:C treatment significantly increased expression of both *cd11b* and *gfap*, although this appeared to be true only at the 24-hour time point (Figures 3A2 and 3B2; *cd11b* main effect of Treatment: *F*(4, 42) = 12.96, *p* < 0.001, η^2^*_p_* = 0.552; *gfap* main effect of Treatment: *F*(4, 42) = 12.992, *p* < 0.001, η^2^*_p_* = 0.553). Sex did not affect the response of either *cd11b* or *gfap* to poly I:C (insignificant Sex x Treatment interactions: *cd11b*: *F*(4, 42) = 0.684, *p* = 0.607; *gfap*: *F*(4, 42) = 0.923, *p* = 0.460).

#### 3.3.2. Interleukins

Poly I:C caused increased expression of *il-1α*, *il-1β*, and *il-6* in both males and females (Figures 3C2, 3D2, and 3E2; main effects of Treatment: *il-1α*: *F*(4, 42) = 9.784, *p* < 0.001, η^2^*_p_* = 0.482; *il-1β*: *F*(4, 42) = 9.512, *p* < 0.001, η^2^*_p_* = 0.475; *il-6*: *F*(4, 42) = 22.28, *p* < 0.001, η^2^*_p_* = 0.680). In males, expression began to increase at the 2-hour time point following poly I:C treatment for *il-1α* (*p* = 0.015), *il-1β* (*p* = 0.057), and *il-6* (p = 0.029), showed peaks at the 4- hour time point (*p* < 0.001 for all), and decreased to saline-treated levels by 24 hours (*p* = 1.00 for all). Each of these genes also showed an overall greater expression in males than females (main effects of Sex: *il-1α*: *F*(1, 42) = 6.398, *p* = 0.015, η^2^*_p_* = 0.132; *il-1β*: *F*(1,42) = 6.695, *p* = 0.013, η^2^*_p_* = 0.137; *il-6*: *F*(1, 42) = 21.1, *p* < 0.001, η^2^*_p_* = 0.334), and a significantly greater magnitude of response in males compared with females (Sex x Treatment interactions: *il-1α*: *F*(4, 42) = 3.103, *p* = 0.025, η^2^*_p_* = 0.228; *il-1β*: *F*(4, 42) = 4.288, *p* = 0.005, η^2^*_p_* = 0.290; *il-6*: *F*(4, 42) = 15, *p* < 0.001, η^2^*_p_* = 0.588). Post-hoc tests revealed for all three genes, males exhibited an even greater response at only the 4-hour time point compared with females (*p* < 0.05 for all). Notably, the peak *il-1α* and *il-1β* expression in males was roughly 3-fold higher than that of the peak female expression for these cytokines, and the *il-6* peak expression in males was more than 10-fold higher than that of females (Figures 3C2, 3D2, and 3E2).

Males showed greater *il-10* gene expression across all time points compared with females (Figure 3F2; main effect of Sex: *F*(1, 39) = 25.642, *p* < 0.001, η^2^*_p_* = 0.397). Additionally, poly I:C significantly increased gene expression of *il-10* in males, but not females (Sex x Treatment interaction: *F*(4, 39) = 3.304, *p* = 0.02, η^2^*_p_* = 0.253). Specifically, male expression of *il-10* at the 4-hour time point following poly I:C was significantly greater than that of saline-treated controls (*p* = 0.001), and this was also greater than the 4-hour expression in females (*p* = 0.001).

#### 3.3.3. Tumor Necrosis Factor Alpha

Gene expression of *tnfα* increased in response to poly I:C, males had significantly higher expression than females overall, and males showed a greater magnitude of response compared with females (Figure 3J2; main effect of Treatment: *F*(4, 42) = 6.407, *p* = 0.0004, η^2^*_p_* = 0.379; main effect of Sex: *F*(1, 42) = 10.1, *p* = 0.003, η^2^*_p_* = 0.194; Sex x Treatment interaction: *F*(4, 42) = 4.117, *p* = 0.007, η^2^*_p_* = 0.282). Post-hoc tests showed that males 4 hours post-treatment had significantly greater expression than those treated with saline (*p* < 0.001), and this was again greater than the 4-hour peak expression in females (*p* = 0.001).

#### 3.3.4. Interferons

Both *ifnα* and *ifnγ* showed a similar response pattern to poly I:C, whereby males treated with poly I:C exhibited a significant acute increase in gene expression of both cytokines, but females did not show the same response (Figures 3G2 and 3I2; *ifnα*: main effect of Treatment: *F*(4, 42) = 5.007, *p* = 0.002, η^2^*_p_* = 0.323; Sex x Treatment interaction: *F*(4, 42) = 3.35, *p* = 0.018, η^2^*_p_* = 0.242; *ifnγ*: main effect of Treatment: *F*(4, 40) = 4.698, *p* = 0.003, η^2^*_p_* = 0.32; Sex x Treatment interaction: *F*(4, 40) = 4.178, *p* = 0.006, η^2^*_p_* = 0.295). Specifically, 4 hours after poly I:C treatment, males showed significantly elevated expression compared to the saline-treated controls (*p* = 0.0001), and this was greater in magnitude than the 4-hour time point in females (*p* = 0.001). Female *ifnα* and *ifnγ* did not seem to respond to poly I:C treatment at any time point.

In contrast, poly I:C transiently increased expression of *ifnβ* in both males and females, and there were no sex differences seen in magnitude of expression increase (Figure 3H2; main effect of Treatment: *F*(4, 42) = 4.855, *p* = 0.003, η^2^*_p_* = 0.316; insignificant Sex x Treatment interaction: *F*(4, 42) = 1.297, *p* = 0.287). Unlike all other cytokines examined in this study, peak expression appeared to be at the 2-hour time point, and expression began decreasing again by 4 hours post-treatment. The magnitude increase was also notable, with a 100-fold increase in females and a 300-fold increase in males.

#### 3.3.5. Chemokines

Poly I:C significantly increased the expression of both *ccl2* and *cxcl10* in males and females, with a peak increase in expression at 4-hours post-infusion (Figures 3K2 and 3L2; main effects of Treatment: *ccl2*: *F*(4, 41) = 25.47, *p* < 0.001, η^2^*_p_* = 0.713; *cxcl10*: *F*(4, 42) = 87.37, *p* < 0.001, η^2^*_p_* = 0.893). Also consistent with many other cytokines, expression of both *ccl2* and *cxcl10* was greater overall in males compared with females (main effects of Sex: *ccl2*: *F*(1, 41) = 44.55, *p* < 0.001, η^2^*_p_* = 0.521; *cxcl10*: *F*(1, 42) = 92.79, *p* < 0.001, η^2^*_p_* = 0.688). Finally, males showed a markedly greater magnitude of response than did females for both chemokines (Sex x Treatment interactions: *ccl2*: *F*(4, 41) = 20.96, *p* < 0.001, η^2^*_p_* = 0.672; *cxcl10*: *F*(4, 42) = 60.51, *p* < 0.001, η^2^*_p_* = 0.852). Remarkably, male *ccl2* expression peaked at nearly 450-fold greater than the expression of saline-treated males compared to a roughly 20-fold increased peak in females (Figure 3K2). Similarly, *cxcl10* expression in males peaked at nearly 500-times that of saline-treated males while female *cxcl10* expression peaked at just over 40-times greater than saline-treated females (Figure 3L2). These massive increases in gene expression are reflected in the strong effect sizes noted for the interaction effect above. Post-hoc tests confirmed that the male 2- and 4-hour time points post-treatment showed significantly greater gene expression of both *ccl2* and *cxcl10* than saline-treated males (*p* < 0.001). Additionally, both the male 2- and 4-hour time points of both genes proved to be significantly greater than the 2- and 4-hour time points in females, respectively (*p* < 0.01).

### 3.4. Central poly I:C increases select hippocampal cytokine protein levels in males and females

#### 3.4.1. Interleukins

We analyzed several members of the interleukin family of cytokines, including classically inflammatory IL-1*α*, IL-1*β*, IL-2, IL-4, and IL-6, and classically anti-inflammatory IL-10. Of these, IL-1*α*, IL-1*β*, IL-4, and IL-6 significantly increased following ICV poly I:C administration in both males and females (Figures 4A, 4C, 4D, 4E; main effects of Treatment: IL-1*α*: *F*(4, 51) = 3.523, *p* = 0.013, η^2^*_p_* = 0.216; IL-1*β*: *F*(4, 51) = 5.721, *p* = 0.001, η^2^*_p_* = 0.31; IL-4: *F*(4, 51) = 5.146, *p* = 0.001, η^2^*_p_* = 0.288; IL-6: *F*(4,51) = 10.298, *p* < 0.001, η^2^*_p_* = 0.447). In all cases, protein levels appear to increase to a peak 4 hours following poly I:C, similar to the effects seen in mRNA expression. In addition, both IL-1*β* and IL-4 also exhibited a main effect of Sex such that protein levels of these cytokines, regardless of time point, were significantly higher in females compared with males (Figures 4C and 4D; IL-1*β*: *F*(1, 51) = 114.226, *p* < 0.001, η^2^*_p_* = 0.691; IL- 4: *F*(1, 51) = 11.03, *p* = 0.002, η^2^*_p_* = 0.178). No interactions of sex and treatment were found for any of the interleukin cytokines examined here (IL-1*α*: *F*(4, 51) = 0.446, *p* = 0.775; IL-1*β*: *F*(4, 51) = 0.513, *p* = 0.726; IL-4: *F*(4, 51) = 0.982, *p* = 0.426; IL-6: *F*(4, 51) = 1.779, *p* = 0.148). Neither IL-2 nor IL-10 showed any effects of poly I:C treatment in either sex (Figures 4B and 4F; insignificant main effects of Treatment: IL-2: *F*(4, 51) = 1.498, *p* = 0.217; IL-10: *F*(4, 51) = 1.122, *p* = 0.357). Similar to IL-1*β* and IL-4, though, we found that females had overall higher levels of IL-10 than did males (main effect of Sex; *F*(1, 51) = 20.27, *p* < 0.001, η^2^*_p_* = 0.284).

#### 3.4.2. Interferon

We also measured IFN*γ* levels in response to poly I:C. Unlike mRNA expression, IFN*γ* protein levels did not change following poly I:C administration in either sex (Figure 4G; insignificant main effect of Treatment: *F*(4, 52) = 1.93, *p* = 0.119). However, IFN*γ* levels were higher in females relative to males (main effect of Sex: *F*(1, 52) = 150.64, *p* < 0.001; η^2^*_p_* = 0.743). This was consistent with mRNA expression data where saline-treated females also showed significantly higher expression of *ifnγ* at baseline than did males (see Figure 3Q).

#### 3.4.3. Chemokines

Finally, we measured protein levels of the chemokines CCL2 and CXCL10 in both sexes. Both chemokines were significantly affected by central administration of poly I:C and in different ways in males and females (Figures 4H and 4I; CCL2: main effect of Treatment: *F*(4, 46) = 18.517, *p* < 0.001, η^2^*_p_* = 0.617; Sex x Treatment interaction: *F*(4, 46) = 3.381, *p* = 0.017, η^2^*_p_* = 0.227; CXCL10: main effect of Treatment *F*(4, 52) = 14.54, *p* < 0.001, η^2^*_p_* = 0.528; Sex x Treatment interaction: *F*(4, 52) = 2.796, *p* = 0.035, η^2^*_p_* = 0.177). In males, CCL2 levels increased earlier (at 2 hours) post-infusion relative to females. For CXCL10, females took longer for protein levels to begin to decrease as compared to the time course in males, with females still showing the massive elevation at 24 hours post-infusion as they did at 4 hours.

Notably, both CCL2 and CXCL10 levels showed the most substantial increases out of all cytokines measured in protein analysis in the hippocampus. Specifically, female CCL2 levels induced by poly I:C peaked at approximately 4 times that of the saline-treated animals, and male CCL2 levels peaked nearly 8 times that of saline-treated males (Figure 4H). More remarkably, female CXCL10 levels rose roughly 16-fold by 4 hours after poly I:C administration, and males showed a 12-fold increase by 4-hours (Figure 4I).

### 3.5. Summary of mRNA and protein data

Overall, hippocampal mRNA expression and protein levels of most of the cytokines and chemokines examined in this experiment responded to central administration of poly I:C in both males and females. We found significant sex differences in baseline mRNA expression and protein levels of several cytokines, where females showed greater basal levels than males. In addition, we found the magnitude of mRNA expression increases was greater in males than females. Protein data showed this to be true only for 2 chemokines, CCL2 and CXCL10.

The heatmaps shown in Figure 5 indicate that most of the immune signaling molecules affected in the immediate phase following poly I:C treatment peaked at 4 hours for both mRNA expression (Figure 5A) and protein levels (Figure 5B) and returned to levels of saline-treated animals by 24 hours post-infusion.

**Figure 5.**
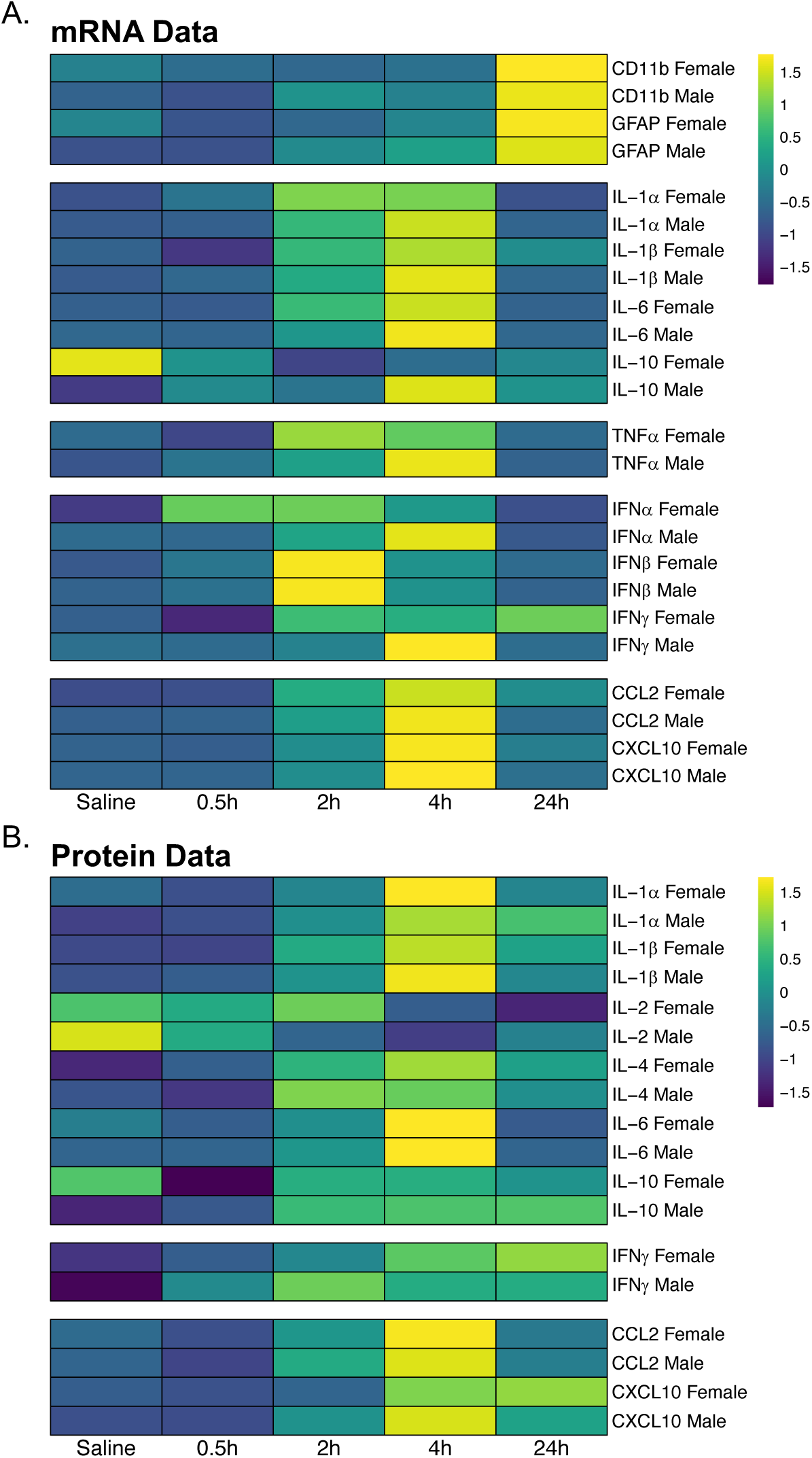
Heatmaps of gene expression and protein levels in the hippocampus. Changes in **(A)** mRNA gene expression and **(B)** protein levels for cytokines, chemokines, and markers of glial activation are shown. Values are centered and scaled across rows to highlight changes across the time course for each gene of interest; thus, differences in magnitude between genes are not depicted.

## 4. DISCUSSION

Here we demonstrated that central administration of poly I:C induced both physiological measures of sickness and concomitant neuroinflammatory activation in both male and female mice. Both males and females treated with poly I:C mounted an acute fever response and showed weight loss that recovered to saline-treated levels 48 hours later. Additionally, both sexes showed acute cytokine and chemokine responses in the hippocampus, as measured by both mRNA expression and protein levels, that followed the time course of fever. Interestingly, mRNA gene expression of *il-1α*, *il-1β*, *il-6*, *il-10*, *ifnα*, *tnfα*, *ccl2*, and *cxcl10* and protein levels of CCL2 and CXCL10 showed a stronger response in males compared with females. Further, gene expression of *il-10*, *ifnα*, and *ifnγ* increased in males only. Given the specific role of type I interferons in viral response (58–60), this sex difference may have important implications for how neuroimmune activation differentially disrupts cognition in males and females.

Poly I:C treatment in both sexes resulted in a significant and transient increase in gene expression and protein levels of most, but not all, cytokines and chemokines in the hippocampus, including IFN*β*, IL-1*α*, IL-1*β*, IL-6, TNF*α*, CCL2, and CXCL10. That administration of an immune stimulant, including viral mimics such as poly I:C, induces a neuroimmune response is not new; however, most of the previous studies on poly I:C used peripheral administration (61–64). As such, multiple, indirect mechanisms are likely involved in causing inflammation in the brain (36). Peripheral administration of poly I:C, specifically, was found to induce neuroinflammation through a separate and distinct pathway than central administration (43). Additionally, these studies examined a subset of cytokines or chemokines, including the most commonly examined IL-1*β*, IL-6, and TNF*α*, and type I interferons that typically respond to viruses. Given evidence of mechanistic complexities governing neuroinflammation, particularly from stimulants such as poly I:C, and given that there are over 300 cytokines with important roles in the immune system and neural function, it is critical to begin looking beyond IL-1*β*, IL-6, and TNF*α* and more strongly consider implications of such limits in experimental design for the field of psychoneuroimmunology.

In addition to the intricacies of how immune stimulants such as poly I:C activate the neuroimmune system, males and females differ in immune and neuroimmune responses. The direction of these differences depends on whether you are looking in the periphery (7) or the brain (26, 27). We found that mRNA gene expression of *il-1α*, *il-1β*, *il-6*, *il-10*, *ifnα*, *tnfα*, *ccl2*, and *cxcl10* and protein levels of CCL2 and CXCL10 in the hippocampus showed a stronger response in males compared with females. A greater magnitude of cytokine and chemokine response in males is consistent with previous findings that male-derived astrocytes have a greater reaction to inflammatory insults compared with females (26,27,65,66). Poly I:C is recognized by microglia, astrocytes, and neurons via toll-like receptor 3 (TLR3) (67–69). The interaction of these 3 cell types is crucial in mediating inflammatory responses (70, 71). Given that TLR3 shows much greater expression in astrocytes relative to microglia (72), the reaction of astrocytes in males may be driving the sex differences in magnitude gene expression response of cytokines following poly I:C. The astrocyte activation marker, GFAP, and the microglial activation marker, CD11b, did not increase until 24 hours after poly I:C treatment and did not show sex differences. However, this does not absolve astrocytes or microglia from the acute response to poly I:C. Specifically, Norden and colleagues found that cytokine gene expression from both astrocytes and microglia preceded increases in astrocyte and microglial activation markers, (GFAP and Iba1, respectively), and that these activation markers similarly did not show reliable increases until the 24-hour time point (73). It is likely, therefore, that a greater astrocytic reaction in males is driving the greater magnitude of response to poly I:C by males.

Viral infections are not immune to observations of sex differences. Specifically, males and females show sex differences in prevalence, intensity, and survival outcomes of viruses (7,8,33). Many report that when females show better survival outcomes or seem more protected against certain viruses, it is due to the female immune system having a greater anti-viral response that allows the virus to be cleared faster (33,74–76). Data on SARS-CoV-2, the virus that causes the coronavirus disease 2019 (COVID-19), has shown a similar sex difference in intensity of infection, with more males having worse symptoms, including neurological complications, and survival outcomes relative to females (77–81). However, in contrast to the data that suggests a greater immune response results in faster recovery in females, recent studies indicate that higher levels of inflammatory cytokines including IL-6, IL-1*β*, CXCL10, and CCL2 are actually correlated with worse COVID-19 outcomes (35, 82). Unequivocally, sex as a biological factor matters in the context of immune and neuroimmune responses as well as in the acute and chronic implications of neuroimmune modulation of neural function, cognition, and emotion in males and females. Limiting studies by only using one sex and extrapolating the data to the other is not only inappropriate given the intricacies of sex differences in the literature discussed here, but it is also potentially dangerous for both development and administration of treatments in inflammatory diseases and disorders.

We observed that most cytokines examined here showed a sex difference in response to poly I:C. Whereas others have reported increases in select inflammatory markers following poly I:C treatment, these studies used either only used male (62, 63) or female rodents (61, 64). To the best of our knowledge, this is the first direct comparison of hippocampal cytokines in males and females as a consequence of poly I:C. Whether the greater magnitude in male response to poly I:C indicates greater neuroprotection or vulnerability is yet to be determined. This is an especially intriguing question given that we also directly compared mRNA expression and protein levels of cytokines and chemokines at baseline and found that females had greater basal immune activity relative to males.

Perhaps it is not only that females are able to mount a greater immune response to help clear viral loads faster and aid in a faster recovery as previously discussed (33,74–76), but also that females start out with greater immune activity that allows them to reach necessary activation states faster than males. Specifically, maybe females do not need to have as great of a magnitude increase in cytokines and chemokines when the neuroimmune system is activated because they already have “more players in the game”. Future work will need to address whether and how sex differences in the cytokine and chemokine basal levels or activation in response to immune challenge result in functional differences in modulation of neural function and contribute to sex-biases in neurological and psychiatric disease.

Of particular note, we observed a sex-specific pattern of expression of the interferon family of cytokines in the hippocampus. Specifically, males showed increases in IFN*α*, IFN*β*, and IFN*γ*, but females only showed a significant response in IFN*β*. This is consistent with previous findings that showed increased gene expression of IFN*β*, but not IFN*α*, in females in response to peripheral poly I:C, though this study did not measure these effects in males for comparison (64). Type I interferons, IFN*α* and IFN*β*, are key to the anti-viral response of the immune system and, as such, are known to respond to viral stimulants including poly I:C (58–60,83,84). Consistent with our data, in which IFN*β* showed an early peak expression levels, type I interferon activity is responsible for inducing inflammatory cytokines such as IL-6 and TNF*α* (62, 64). Additionally, interferon signaling from poly I:C treatment also results in altered glutamatergic signaling (83, 84), which is known to be critical for hippocampal memory formation (85). Given that males induce both IFN*α* and IFN*β* in the hippocampus following poly I:C whereas females only induce IFN*β*, interferons have important implications for sex differences in neuroimmune disruption of cognition from virus or viral-like stimulants.

## 5. CONCLUSION

This study characterizes the sickness and neuroimmune responses to central administration of poly I:C, and we observed sex-specific patterns of hippocampal cytokine responses. Specifically, we identified type I interferons as one potential node mediating sex-specific cytokine responses and neuroimmune effects on synaptic plasticity and cognition. Additionally, the magnitude of response of cytokines such as CCL2 and CXCL10 highlight the importance of future work incorporating a more comprehensive set of inflammatory markers using multiple endpoints. Neuroimmune activation is known to play a role in cognitive deficits and affective dysregulation in diseases such as Alzheimer’s Disease and other dementias (86), Post-Traumatic Stress Disorder (87, 88), depression (4, 89), and now also COVID-19 (90). Given the sex/gender biases in prevalence, severity, and/or survival outcomes, identifying sex-specific neuroimmune responses will provide novel targets for personalized prevention and treatment of these diseases.

## LIST OF ABBREVIATIONS

ANOVA: analysis of variance
CCL2: C-C motif chemokine ligand 2
Cd11b: cluster of differentiation molecule 11B
cDNA: complementary deoxyribonucleic acid
COVID-19: coronavirus disease-19
CXCL10: C-X-C motif chemokine ligand 10
DNA: deoxyribonucleic acid
GAPDH: glyceraldehyde-3-phosphate dehydrogenase
GFAP: glial fibrillary acidic protein
HPRT1: hypoxanthine phosphoribosyltransferase 1
Iba1: ionized calcium-binding adaptor molecule 1
ICV: intracerebroventricular
IFN*α*: interferon *α*
IFN*β*: interferon *β*
IFN*γ*: interferon *γ*
IL-1*α*: interleukin-1*α*
IL-1*β*: interleukin-1*β*
IL-2: interleukin-2
IL-4: interleukin-4
IL-6: interleukin-6
IL-10: interleukin-10
kg: kilogram
LPS: lipopolysaccharide
mg: milligram
mm: millimeter
mRNA: messenger ribonucleic acid
PCR: polymerase chain reaction
Poly I:C: polyinosinic:polycytidylic acid
qPCR: quantitative real-time polymerase chain reaction
RNA: ribonucleic acid
RPLP0: ribosomal protein lateral stalk subunit P0
SARS-CoV-2: severe acute respiratory syndrome coronavirus 2
TLR: toll-like receptor
TNF*α*: tumor necrosis factor *α*
μg: microgram
μL: microliter
μm: micrometer

## DECLARATIONS

### Ethics Approval

All animal protocols used in these experiments were approved by the Institutional Animal Care and Use Committee (IACUC) at the University of Michigan.

### Consent for Publication

Not applicable.

### Availability of Data and Materials

The data used and analyzed for the current study are available from the corresponding author upon reasonable request.

### Author’s Contributions

CKP designed the experiments, performed all animal surgeries and treatments, collected physiological measurements (body temperature and weights) and tissue, processed tissue samples and conducted all molecular biology assays (qPCR and multiplex assays), ran statistical analyses, and wrote the manuscript. RGH aided in the collection of physiological measurements (body temperature and weights) and monitored animals for sickness behaviors and surgical recovery throughout the experiments. NCT provided intellectual contributions regarding experimental design and data interpretation, supplied financial support and laboratory space and equipment to complete the experiments, and contributed significantly to the editing of this manuscript. All authors read and approved the final version of this manuscript.

## Acknowledgements

We would like to thank Dr. Ashley Keiser, Dr. Daria Tchessalova, Brynne Raines, and Sarah Jacob for assistance with the techniques, reagents, and data collection for these experiments.

## Author’s Information

CKP is trained in behavioral neuroscience with specific interests in behavioral neuroimmunology. Since 2013, CKP has worked on models of neuroimmune activation in both rats and mice with the goal of understanding how the neuroimmune system functions to modulate cognition and affect and both whether and how biological sex interacts with these processes.

NCT studies the mechanisms of learning and memory in males and females, the modulation of memory and affective processes by neuroimmune activation, and long-term impact of illness on the hippocampus, cognitive decline, Alzheimer’s Disease, and neuropsychiatric disorders.

